# Precise annotation of human, chimpanzee, rhesus macaque and mouse mitochondrial genomes using 5’ and 3’ end small RNAs

**DOI:** 10.1101/706093

**Authors:** Zhi Cheng, Haishuo Ji, Xiufeng Jin, Bo Wang, Tungon Yau, Ze Chen, Defu Chen, Wenjun Bu, Daqing Sun, Shan Gao

## Abstract

Using 5’ and 3’ end small RNAs, we annotated human, chimpanzee, rhesus macaque and mouse mitochondrial genomes at 1 base-pair (bp) resolution to cover both strands of the mammalian mitochondrial genome entirely without leaving any gaps or overlaps. The precise annotation of all coding and non-coding genes (e.g. ncMT1, MDL2 and MDL1AS) led to the discovery of novel functions and mechanisms of mitochondrion. In this study, we defined the conserved sequence block (CSB) region to span five CSBs (CSB1, CSB2, CSB3, LSP and HSP) and identified the motifs of five CSBs in the mitochondrial displacement loop (D-loop) regions of 52 mammals. The conserved arrangement of these five CSBs in 17 primates inspired us to investigate the function of the mtDNA D-loop, which has been puzzling scientists for more than 50 years. We found that 5’ sRNAs of MDL1AS control the expression levels of mitochondrial genes as a whole by a negative feedback mechanism. Thus, the precise annotations of three CSBs (CSB2, LSP and HSP) in more species will help to understand the function of the mtDNA D-loop. The precision annotation of animal mitochondrial genomes also provides abundant information for studying the molecular phylogenetics and evolution of animals.

## Introduction

The annotation of animal mitochondrial genomes provides information for the study of the molecular phylogenetics and evolution of animals and for the investigation of RNA processing, maturation, degradation and gene expression regulation [1]. Although the annotation of mitochondrial mRNAs, tRNAs, and rRNAs can be easily performed using web servers (*e.g.* MITOS [2]), the current methods (e.g. Blastx or structure-based covariance models) result in gaps and overlaps in the annotations of mitochondrial genomes and the annotation resolution is limited. In 2017, we constructed the first quantitative transcription map of animal mitochondrial genomes by sequencing the full-length transcriptome of the insect *Erthesina fullo* Thunberg [3] on the PacBio platform [4] and established a straightforward and concise methodology to improve genome annotation. In 2019, we used 5’ and 3’ end small RNAs (5’ and 3’ sRNAs) [1] to improve genome annotation at 1 bp resolution [5], which was later named as precise annotation.

In this study, we performed precise annotation of human, chimpanzee, rhesus macaque and mouse mitochondrial genomes to cover both strands of the mammalian mitochondrial genome entirely of all coding and non-coding genes without leaving any gaps or overlaps. We identified the H-strand promoters (HSPs) and the L-strand promoters (LSPs) of these mitochondrial genomes precisely. Furthermore, we defined the conserved sequence block (CSB) regions to span five CSBs (CSB1, CSB2, CSB3, LSP and HSP) and investigated their motifs and arrangement in 52 mammalian mitochondrial genomes. To demonstrate significance of the precise annotation method that we used here to annotate mitochondrial genomes, we investigated the function of the mtDNA D-loop, which has puzzled scientists for more than 50 years. We found that 5’ sRNAs of MDL1AS control the expression levels of mitochondrial genes as a whole by a negative feedback mechanism.

## Results

### Improvement of precise annotation

Based on 5’ and 3’ end small RNAs (5’ and 3’ sRNAs), the *Homo sapiens*, *Pan troglodytes*, *Macaca mulatta* and *Mus musculus* mitochondrial genomes were annotated precisely (**Supplementary file 1**) to update the previous version of annotations (NCBI RefSeq: NC_012920.1, NC_001644.1, NC_005943.1 and NC_005089.1), respectively. The new annotations of the human and mouse mitochondrial genomes were confirmed by the full-length transcriptome data (**Materials and Methods**). The two strands of one mitochondrial genome are differentiated by their nucleotide content. They are a guanine-rich (G-rich) strand referred to as the Heavy strand (H-strand) and a cytosine-rich strand referred to as the Light strand (L-strand). The precise annotations generated in this study covered the entire H-strand and L-strand of the mammalian mitochondrial genome without leaving any gaps or overlaps. The new annotations of the four mitochondrial genomes confirmed our previous findings related to the human mitochondrial genome [6]. Particularly, it was confirmed that two polycistronic transcripts, COI/tRNA^Ser^AS and ND5/ND6AS/tRNA^Glu^AS, are not further cleaved but used as templates for protein synthesis. The new annotation of COI/tRNA^Ser^AS modified the ‘mitochondrial cleavage’ model that we proposed earlier [5]. According to the new model, RNA is not cleaved between mRNAs and their downstream antisense tRNAs. Long antisense genes (usually above 1000 bp) in four mammalian mitochondrial genomes had to be predicted and then annotated using this new ‘mitochondrial cleavage’ model, as they are transcribed as transient RNAs (e.g. tRNA^Met^AS/ND2AS/tRNA^Trp^AS) and are usually not well covered by aligned reads from sRNA-seq or RNA-seq data owing to their rapid degradation. Two lncRNAs (MDL2 and MDL1AS) and one sRNA non-coding mitochondrial RNA 1 (ncMT1) was identified as a functional ncRNA in mammalian mitochondrial genomes, while all other reported mitochondrial ncRNAs (e.g. lncND5 and lncND6 [7]) were degraded fragments of transient RNAs or random breaks that occurred during experimental processing. In the new annotations, long antisense genes were annotated as H-strand Antisense Segments (HASn) or L-strand Antisense Segments (LASn), where n starts counting from behind the D-loop region. The results from this study did not support that the previously reported lncND5, lncND6, and lncCytb [7] are lncRNAs, as they are not likely to have specific functions as transient RNAs. Using our ‘mitochondrial cleavage’ model, lncCytb was annotated as LAS3 (CytbAS/tRNA^Thr^AS) in the new annotations, whereas lncND6 and lncND5 were degraded fragments of two other annotated transient RNAs (ND5/ND6AS/tRNA^Glu^AS and LAS2) respectively (**Supplementary file 1**).

The precise annotations corrected the previous annotations of mitochondrial coding genes (mRNA genes) by the analysis of their Open Reading Frames (ORFs). The mistakes with the previous annotations of mitochondrial coding genes were usually attributed to the irregularity of the start codons (e.g. TTG in *Erthesina fullo* [3]) and stop codons (e.g. TA and T) in their Coding Sequences (CDSs), which usually begin at the 5’ ends by start codons ATG, ATA or ATT and end at the 3’ ends by stop codons TAA, TA and T. However, TA and T are completed by polyadenylation to form TAA after RNA cleavage and are thus often ignored. If one A/G or AA/AG is downstream of TA (e.g. ATP6) or T (e.g. ND1 and ND2), TAA or TAG will be wrongly annotated as the stop codon. By using the precise annotation method as in this study, one coding gene is annotated by its mature RNA rather than its CDS, which could result in several nucleotides being annotated before the start codon in the CDS. One typical example is the three nucleotides before ATG of COI in mammals. These three nucleotides CTT, CTG, CTT, CTC, CTA and CAT appear 32, 5, 5, 4, 3, 2 and 1 times in 52 mammalian mitochondrial genomes respectively (**Supplementary file 1**). These three nucleotides are not likely to belong to start codons as one, two or more than three nucleotides appear frequently before start codons of other mitochondrial coding genes in mammals. For example, ACATA(start codons underlined), ACACACCCATG, CCATG and GTGTTCTTTATT were discovered at the beginning of ND1 in human, chimpanzee, rhesus macaque and mouse mitochondrial genomes respectively. ATC is often identified as a start codon (e.g. ATCAATATT in mouse ND5) for mitochondrial coding genes [8], we suspected if ATC belongs to start codons as it often appears before ATG, ATA or ATT.

In our previous study, we annotated the mitochondrial tRNAs of human at 1 bp resolution and proposed a mitochondrial tRNA processing model [1]. In this model, one mitochondrial tRNA is cleaved from a primary transcript into a precursor, and then the amino acid acceptor stem of the precursor is adenylated (for the blunt ends, e.g. tRNA^Tyr^ in human) or trimmed (for the overhangs longer than 1 bp, e.g. tRNA^Asn^ in human) to contain a 1-bp overhang at the 3’ end. The precise annotations of mitochondrial tRNAs revealed the differences in overhangs of the amino acid acceptor stems (**Figure 1**) between mouse and three primates (human, chimpanzee and rhesus macaque). In our previous study, we also improved the annotations of consecutive tRNAs (e.g. tRNA^Tyr^/tRNA^Cys^/ncMT1/tRNA^Asn^/tRNA^Ala^ in human) [1]. In particular, we found a novel 31-nt ncRNA named non-coding mitochondrial RNA 1 (ncMT1) between tRNA^Cys^ and tRNA^Asn^ (**Figure 2A**). The mature RNA of ncMT1 had polyA at its 3’ end and formed a typical stem-loop structure with 8-bp perfect matched nucleotides in the stem. Among potential orthologs of ncMT1 genes in 52 mammalian mitochondrial genomes (**Supplementary file 1**), 47 orthologs had the identical 8-bp perfect matched nucleotides and five orthologs (**Figure 2B**) had co-varied mutations (C-G/A-T) to maintain a conserved stem-loop structure. This highly evolutionary conservation implied that ncMT1 had specific functions in mammals.

**Figure 1.**
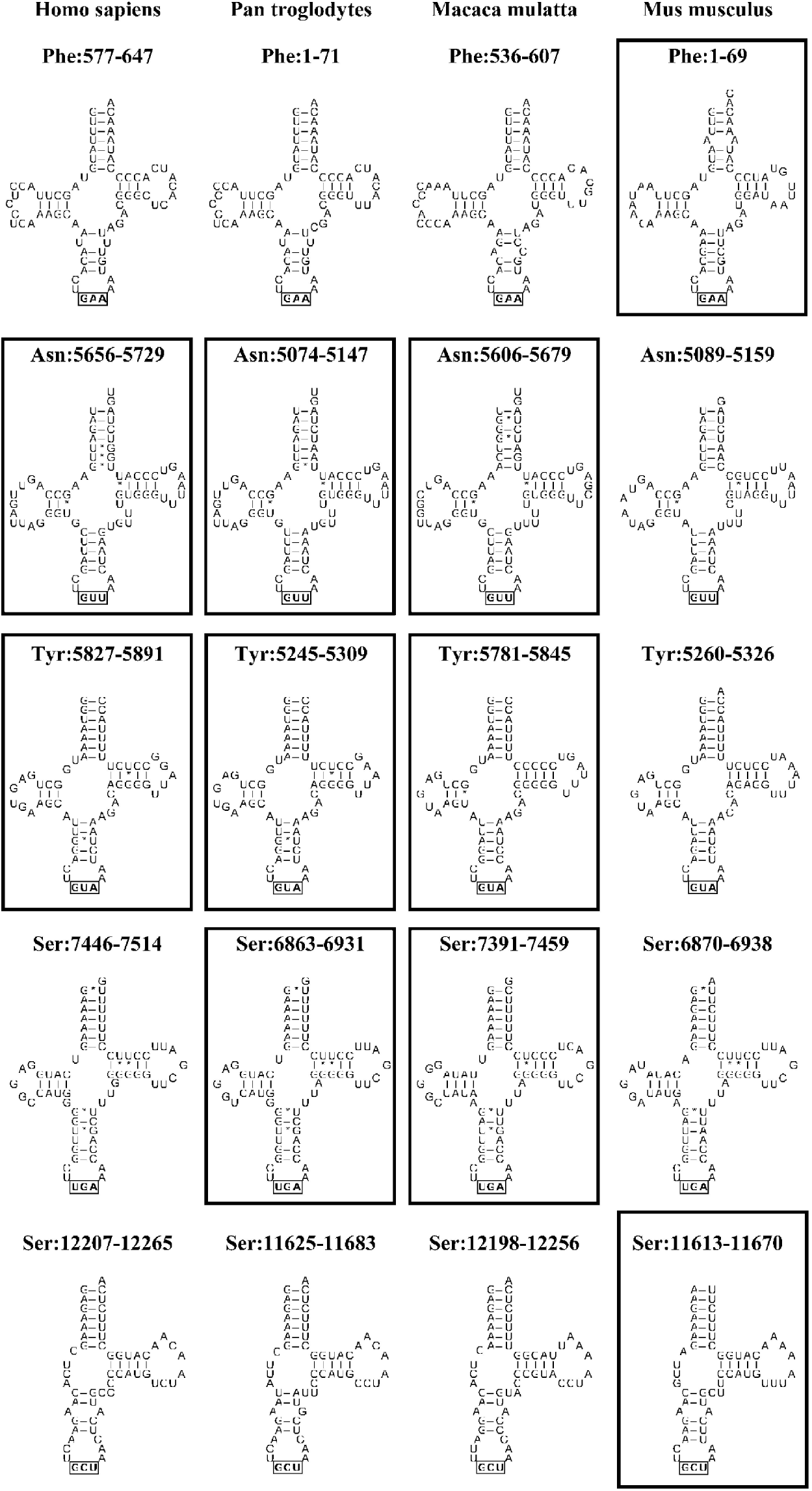
Precise annotation of mitochondrial tRNAs. The corrected annotations of mitochondrial tRNAs are marked in black box. In the precise annotations, mitochondrial tRNAs are annotated as their precursors rather than their mature RNAs. The mouse tRNA^Phe^, tRNA^Asn^, tRNA^Tyr^ and tRNA^Ser(11613-11670)^ have 2, 1, 2 and 0 overhangs in the amino acid acceptor stems, while three primates (human, pan troglodytes and macaca mulatta) have 1,2,0 and 1 overhangs.

**Figure 2.**
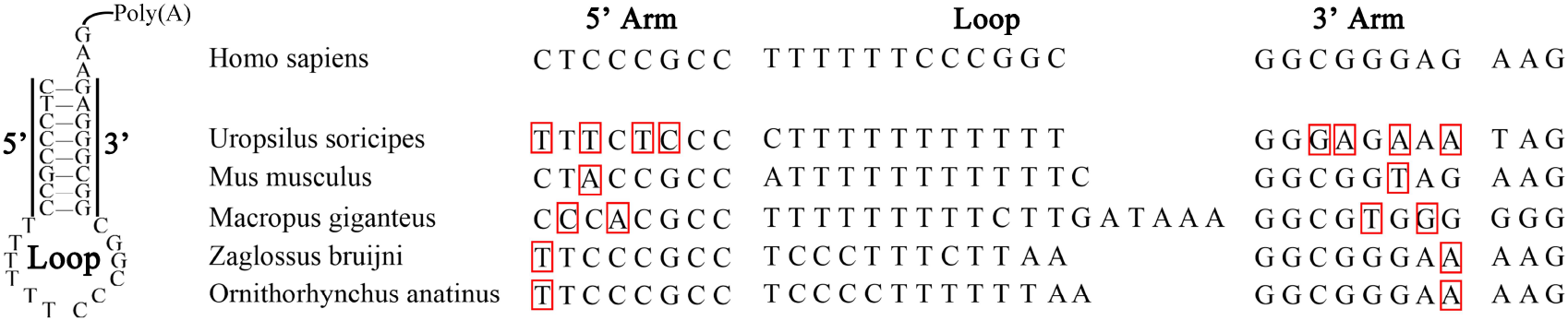
The evolutional conservation in ncMT1. The sRNA ncMT1 was identified as a functional ncRNA in mammalian mitochondrial genomes. Among 52 mammalian mitochondrial genomes, five orthologs have co-varied mutations (C-G/A-T), while others have identical sequences.

### MDL1, MDL2, MDL1AS and MDL2AS

One animal mitochondrial genome contains at least one control region (CR) with a few possible exceptions (e.g. Asymmetron [1]). CR is also called the Displacement-loop (D-loop) region (**Figure 3A**), although the control and the D-loop region is not the same in most species. In our previous study, we proposed that any animal mitochondrial genome that contains one control region transcribes both the H-strand and the L-strands entirely. One control region contains at least two Transcription Initiation Sites (TISs), which are the TIS of the H-strand (TIS_H_) and the TIS of the L- strand (TIS_L_) [5]. TIS_H_ and TIS_L_ belong to two independent regions, the H-strand promoter (HSP) and the L-strand promoter (LSP). Using 5’ and 3’ sRNAs, we were able to precisely determine TIS_H_s and TIS_L_s in human, chimpanzee, rhesus macaque and mouse mitochondrial genomes (**Supplementary file 1**). Particularly, the precise annotation of mouse TIS_H_ and TIS_L_ at the positions 16,285 and 16,188 corrected the previous annotations at the positions 16,295 and 16,187 (corrected using NC_005089.1)[9], respectively. The new annotation of a unique mouse TIS_H_ at the positions 16,285 also corrected the previous annotations of two TIS_H_s at the positions 16,295 and 16,287 (corrected using NC_005089.1) or even more TIS_H_s proposed in the previous study [9].

One CR involves four RNAs (**Figure 3B**). On the H-strand, the shorter RNA is defined as Mitochondrial D-loop 1 (MDL1) and the longer RNA is defined as MDL2. On the L-strand, the shorter RNA is defined as Mitochondrial D-loop 1 antisense gene (MDL1AS) and the longer one is defined as MDL2AS [5]. MDL1 and MDL1AS start at the TIS_H_ and TIS_L,_ and end at the downstream cleavage sites on the H-strand and the L-strand, respectively. MDL2 and MDL2AS reside between two nearest cleavage sites on the H-strand and L-strand, respectively. In our previous study, the longer RNA on the H-strand was defined as the human MDL1 (hsa-MDL1), because the shorter one (NC_012920: 561-576) is only 16 bp in length, which is not likely to have specific functions. In the new annotation of human mitochondrial genome, MDL1 was renamed as MDL2, thus MDL2 and MDL1AS in human were annotated as lncRNAs. While the full-length transcripts of MDL1, MDL2 and MDL1AS have been detected [6], those of MDL2AS have not been detected in human, chimpanzee, rhesus macaque and mouse mitochondrial genomes.

**Figure 3.**
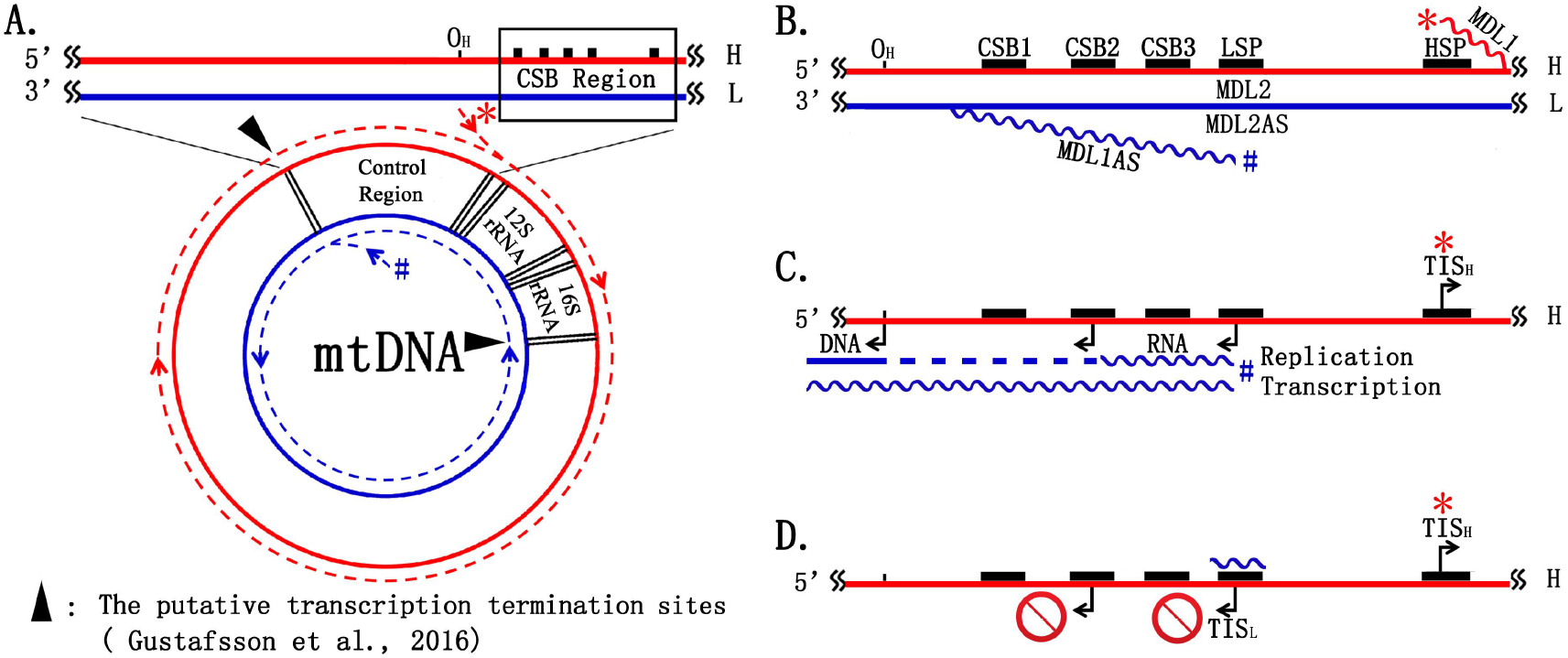
Five CSBs and their arrangement. **#**TIS_H_ and **\***TIS_L_. **A**. Any animal mitochondrial genome that contains one control region transcribes both entire strands. Full-length transcripts of MDL2AS (extending the end of L-strand primary transcript from 16S rRNA to D-loop) have been not detected in human, pan troglodytes, macaca mulatta and mouse mitochondrial genomes. **B**. The conserved arrangement of these five CSBs. **C**. The H-strand replication is intimately linked with the L-strand transcription. **D**. The excessive 5’ sRNAs of MDL1AS could inhibit the L-strand transcription and thus the H-stran dreplication by RNA-DNA hybrid.

### Precise annotation of five conserved sequence blocks

Comparisons of D-loops in human and mouse mtDNAs originally revealed the presence of three Conserved Sequence Blocks (CSBs), referred to as CSB1, -2, and -3 [10] (**Figure 3A**). Compared with CSB2 that is responsible for the transition from RNA to DNA in the H-strand replication, the biological functions of CSB1 and CSB3 are still unclear. CSB3 is the most evolutionarily conserved among the five CSBs, while CSB1, CSB2, LSP and HSP are only conserved within one order of the class Mammalia based on the analytical results using 52 mammalian mitochondrial genomes. Based on precise annotations of CSB1s, CSB2s, CSB3s, LSPs, and HSPs in human, chimpanzee, rhesus macaque and mouse mitochondrial genomes (**Figure 3B**), we predicted CSBs in 48 of 52 mammalian mitochondrial genomes (**Supplementary file 1**). Then, we used all the precisely annotated and predicted CSBs in 4 and 48 mammal, particularly 17 primate mitochondrial genomes to identify the sequence motifs. On the H-strand, the D-loop region has many ployA (A_m_) and ployC (C_n_) patterns allowing single nucleotide polymorphisms (SNPs) or Insertions/Deletions (InDels), which could provide signals for specific functions (e.g. recognition by enzyme for the initiation of DNA replication or transcription). In these patterns, the most frequent SNPs are A/G or C/T. As the most evolutionarily conserved region in 52 mammals, CSB3 has a motif A_3_C_4_A_5_ allowing SNPs or InDels, where A_5_ is the core of this motif. The previous study reported that two closely related 15-nt sequence motifs ATGN_9_CAT on each side of the D-loop [11]. One copy forms the motif of CSB1, whereas the other motif is located upstream in the reference genome. CSB1s in 52 mammalian mitochondrial genomes were identified to share this motif with three exceptions, which are ATAN_9_CAT in *Pteropus vampyrus* and ATGN_8_CAT in *Pongo abelii* and *Pongo pygmaeus*. In 17 primates (**Table 1**), HSP has a motif A_3_C_4_AAAGA, where AAAGA is the core of this motif. The reverse complimentary sequence of LSP has the same core AAAGA as HSP and a similar motif but allows more SNPs or InDels than HSP. CSB2 has a motif A_3_C_6_ allowing more SNPs or InDels than HSP, where C_6_ is the core of this motif. All the motifs in CSB2, CSB3 and HSP contain A_3_C_4_, which proves that A_m_ and C_n_ patterns provide signals for specific functions.

**Table 1.**
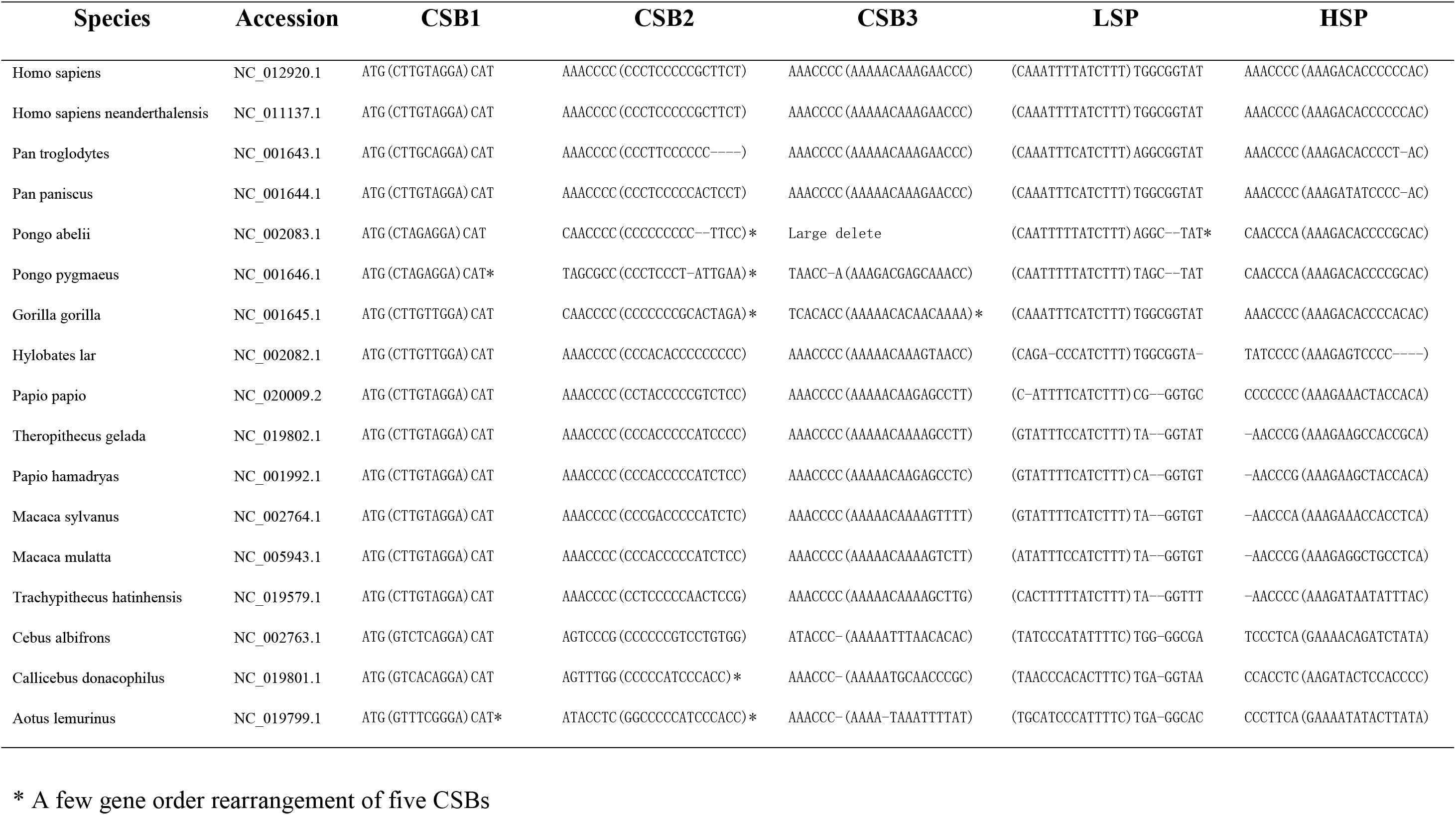
The arrangement of five CSBs

As above results showed that HSP and LSP are also CSBs, we define the CSB region to span CSB1, CSB2, CSB3, LSP and HSP (**Figure 3A**). As one important finding, we found that the arrangement of five CSBs is evolutionarily conserved in 17 primates with several exceptions (**Table 1**). HSP is always close to the 3’ end of the D-loop region and has a distance ranging from 10 to 50 bp to the 3’ end based on our data, although the HSP motif has large sequence diversity between orders or families. The core of HSP motif in primates is AAAGA, while the core of HSP motif in rodents is also A-rich but allows Ts (e.g. AATAA in mouse) and the core of HSP motif in carnivores allows more Ts (e.g. AATTT in *Canis lupus familiaris* or AAATT in *Puma concolor*, *Catopuma temminckii* and *Lynx rufus*). Four other CSBs have steady distances to HSP. Using 17 mitochondrial genomes, we estimated the distance between CSB1 and CSB2, CSB2 and CSB3, CSB3 and LSP, and LSP and HSP to be about 100, 50, 50, and 100–150 bp respectively. The distance between LSP and HSP varies greatly among the 17 primates.

Based on the findings above, we proposed a simple method to identify five CSBs. First, TIS_H_s and TIS_L_s can be identified using 5’ and 3’ sRNAs (**Materials and Methods**). Then, CSB2 can be identified (e.g. for chimpanzee, rhesus macaque or mouse) by the polyC motif at upstream of LSP in the reference genome. The arrangement of five CSBs and the steady distances between them can be used to determine CSB1 and CSB3 (ATGN_9_CAT). As five CSBs are evolutionarily conserved, information from close species can be used to predict five CSBs in one family or order. In this way, we predicted five CSBs in 48 of 52 mammalian mitochondrial genomes (**Supplementary file 1**). Tandem repeats in the D-loop region are obstacles to estimating distances between five CSBs. However, tandem repeats exist in at least 19 of 52 mammalian mitochondrial genomes, particularly in all 12 carnivores. Although the biological functions of tandem repeats in the D-loop region are still largely unknown, those in the *Didelphis virginiana* mitochondrial genome could contain the repeated HSP (**Supplementary file 1**). In one of our previous studies, we found the same phenomenon in the *E. fullo* mitochondrial genome. In the *E. fullo* mitochondrial genome, HSP is also close to 3’ end of the D-loop region as mammals. This suggests that the conserved arrangement of five CSBs or at least three CSBs (CSB2, LSP and HSP) may exist in a wide range of animal mitochondrial genomes.

### 5’ sRNAs of MDL1 and MDL1AS

Although the existence of D-loop has been known for over fifty years, it is still not understood why this structure is synthesised and maintained [12]. The well accepted theory is that D-loop involves in mtDNA replication or transcription, and so indirectly regulates these processes. A long-standing hypothesis is that the H-strand replication is intimately linked with the L-strand transcription. Further studies of D-loop regions in human and mouse mtDNAs have suggested that short mitochondrial transcripts, originating at ITL, serve as primers for the initiation of synthesis of nascent H-strands. Although a series of proteins (e.g. TFAM, TFB2M and TWINKLE [13]) encoded by nuclear genes have been reported to regulate mtDNA replication or transcription in the following studies [8], it had not been considered that mtDNA could regulate its replication or transcription by itself until the discovery of MDL1, MDL1AS and their 5’ sRNAs [6]. In the previous study [6], our hypothesis was that 5’ sRNAs of MDL1 and MDL1AS control the expression levels of mitochondrial genes as a whole by a negative feedback mechanism. In particular, the excessive 5’ sRNAs of MDL1AS could inhibit the L-strand transcription and thus the H-strand replication by RNA-DNA hybrid (**Figure 3CD**). In this study, we performed the preliminary experiments to prove this hypothesis. First, we used *in situ* hybridization to validate that the synthetic DNA/RNA can permeate mitochondrial inner membrane (**Figure 4ABC**). Then we transfected a 37-bp synthetic RNA that mimics the 5’ sRNAs of MDL1AS into HEK293 cells. The results of 3-(4,5-dimethyl-2-thiazolyl)-2,5- diphenyl-2-H-tetrazolium bromide (MTT) experiments (**Materials and Methods**) showed that the viabilities of cells treated by the 37-bp synthetic RNA increased with the increasing concentrations of synthetic RNA, but decreased significantly after ×3 dosages, while the viabilities of cells treated by the reverse complimentary of the 37- bp synthetic RNA did not change significantly (**Figure 4D**). Therefore, 5’ sRNAs of MDL1AS controlled the expression levels of mitochondrial genes as a whole by a negative feedback mechanism.

**Figure 4.**
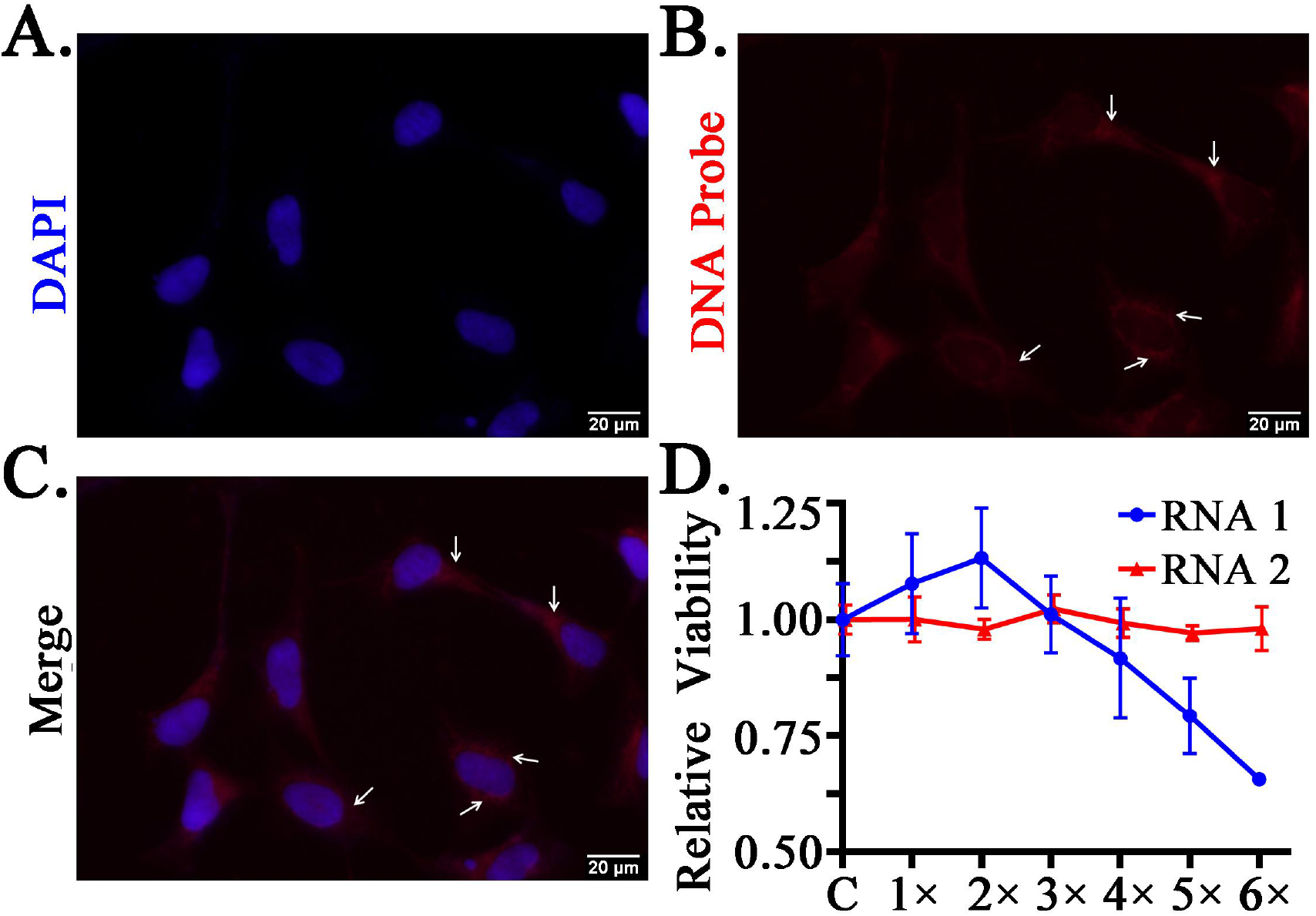
5’ sRNAs of MDL1AS and their functions. A. Cell nuclei (in blue color) were stained with DAPI. **B.**Targets were *in situ* hybridized with biotin-labeled DNA probes stained in red color. **C.**The merged figure showed the mitochondrion (in red color) around the cell nuclei (in blue color). **D.**The relative activities were calculated by the average activity of samples in the treatment group divided by that of samples in the control group. RNA1 and -2 represent 5’ sRNAs of MDL1AS and their reverse complimentary sRNAs.

The precise annotations generated in this study extend the transcriptional regions on the H-strand by the discovery of MDL2 and the L-strand by the sRNA-seq data (**Figure 3A**). However, we did not find the assumed Transcription Termination Sites (TTSs) of two primary transcripts in the D-loop region. In our previous study, we found that RNA polymerase had the ability to read through the assumed TTS after the H-strand primary transcript had been completely synthesized. As cells usually prefer economic and efficient ways for existence, the ‘read through’ model [6] supports uninterrupted transcription of the animal mitochondrial genomes. As the H-strand and the L-strand are transcribed as two primary transcripts, the 5’ sRNAs of MDL1 and MDL1AS can be used to quantify the expression levels of these two primary transcripts, respectively. Using the human931 sRNA-seq dataset, we found that the 5’ sRNAs of MDL1 increased in the cancer samples, while the 5’ sRNAs of MDL1AS decreased in the cancer samples (**Supplementary file 2**). These cancer types included hepatocellular carcinoma (SRA: SRP002272) (**Table 2**), colorectal cancer (SRA: SRP022054), adrenocortical carcinoma (SRA: SRP028291) and head and neck squamous cell carcinoma (ENA: ERP001908). One explanation for the increase of 5’ sRNAs of MDL1 in cancer cells is higher transcription initiation of mitochondrial genes in these cells to produce more MDL1 5’ sRNAs. However, another explanation could be increased occurrence of uninterrupted transcription of animal mitochondrial genomes in normal cells in comparison to cancer cells. Furthermore, the decrease of 5’ sRNAs of MDL1AS in cancers may involve additional factors, which is beyond the scope of this study.

**Table 2.**
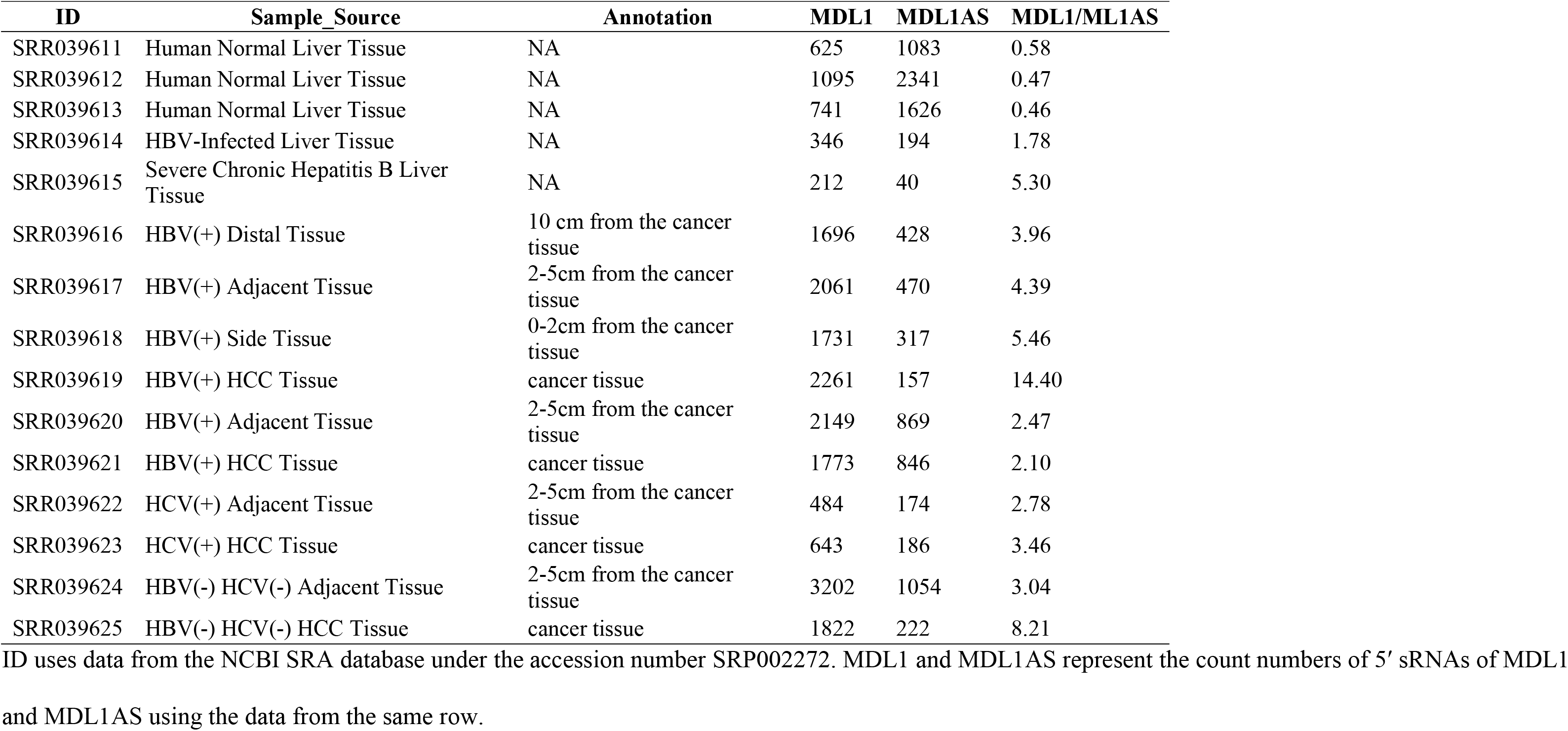
5’ sRNAs of MDL1 and MDL1AS are quantified

## Conclusion and Discussion

In this study, we performed the precise annotation of human, chimpanzee, rhesus macaque and mouse mitochondrial genomes. These annotations provide abundant information for studying the molecular phylogenetics and evolution of animals. We defined the conserved sequence block (CSB) region to span five CSBs (CSB1, CSB2, CSB3, LSP and HSP) and identified the motifs of five CSBs in the mitochondrial displacement loop (D-loop) regions of 52 mammals. The conserved arrangement of five CSBs or at least three CSBs (CSB2, LSP and HSP) may exist in a wide range of animal mitochondrial genomes. As a long-standing hypothesis is that the H-strand replication is intimately linked with the L-strand transcription, CSB2 requires to be downstream LSP for the RNA-DNA transition. Unexpectively, we found a few gene order rearrangement of three CSBs (**Table 1**). In the pongo abelii mitochondrial genome, CSB2 is not downstream LSP. This findings suggests that other C-rich regions can also be responsible for the functions of CSB2. Future studies need be conducted to investigate the arrangement of CSB2, LSP and HSP in a wide range of animal mitochondrial genomes. This will help to eventually reveal mechanisms in the RNA-DNA transition and even the functions of D-loop.

As animals transcribe both entire strands of its mitochondrial genome to produce two primary transcripts, the regulation on the transcription iniation and termination is more worthy of study. Our preliminary studies proved that 5’ sRNAs of MDL1AS controlled the expression levels of mitochondrial genes as a whole by a negative feedback mechanism. however, the underlying mechanisms are still unclear. Using the human931 sRNA-seq dataset, we found that the 5’ sRNAs of MDL1 increased in the cancer samples, while the 5’ sRNAs of MDL1AS decreased in the cancer samples. Future studies need be conducted to investigate the relationships of these changes in 5’ sRNAs of MDL1 or MDL1AS with the regulation of two primary transcripts. This will help to eventually reveal mechanisms of the mitochondrial gene transcription and its regulation on the RNA level.

## Materials and Methods

The reference sequences of human, pan troglodytes, macaca mulatta and mouse mitochondrial genomes were downloaded from the NCBI RefSeq database under the accession numbers NC_012920.1, NC_001644.1, NC_005943.1 and NC_005089.1, respectively. The sRNA-seq data were downloaded from the NCBI SRA database under the accession numbers SRP017691, SRP012018, SRP059657. The mouse full-length transcriptome data was downloaded from the NCBI SRA database under the accession numbers SRP067402 and SRP101446. The human mitochondrial genome was annotated using 5’ and 3’ sRNAs from the human931 sRNA-seq dataset, which had been constructed in our previous study [14]. The mouse mitochondrial genome was annotated using 5’ and 3’ sRNAs from the datasets SRP017691 and SRP012018. The pan troglodytes and macaca mulatta mitochondrial genomes were annotated using 5’ and 3’ sRNAs from the dataset SRP059657.

The precise annotation using 5’ and 3’ sRNAs followed the protocol published before [1]. The precise annotation of human mitochondrial genome was confirmed using pan RNA-seq analysis [1] and The precise annotation of mouse mitochondrial genome was confirmed using the mouse full-length transcriptome from datasets SRP067402 and SRP101446. Data cleaning and quality control were performed using Fastq_clean [15]. Fastq_clean is a Perl based pipeline to clean DNA-seq [16], RNA- seq [17] and sRNA-seq data [14] with quality control and had included some tools to process Pacbio data in the version 2.0. Statistics and plotting were conducted using the software R v2.15.3 with the package ggplot2 [18]. The 5’ and 3’ ends of mature transcripts, polycistronic transcripts, antisense transcripts and the positions of 5’ m7G caps were observed and curated using the software Tablet v1.15.09.01 [19].

One synthetic DNA and one synthetic RNA (RNA1) were used to represent the 5’ sRNAs of MDL1AS. Three other synthetic RNAs (RNA2, -3 and -4) were used to represent the reverse complimentary of RNA1, 5’ sRNAs of MDL1 and the reverse complimentary of RNA3. One synthetic RNA (Control) was used as control for the MTT experiments. The sequences of DNA, RNA1, -2, -3, -4 and Control are 5’ biotin-CCGCCAAAAGATAAAATTTGAAATCTGGTTAGGCTGG, CCGCCAAAAGATAAAATTTGAAATCTGGTTAGGCTGG, CCAGCCTAACCAGATTTCAAATTTTATCTTTTGGCGG, AAACCCCAAAGACACCCCCCACAGT, ACTGTGGGGGGTGTCTTTGGGGTTT and CCTCTTACCTCAGTTACAATTTATA. The *in situ* hybridization experiments using 15 nmol synthetic DNA followed the same procedure as described in one of our previous studies [20]. The MTT experiments using RNA1, -2, -3, -4 and Control followed the same procedure as described in one of our previous studies [21] with a little modification. The modification was that the cells were cultured 24 hours before RNA transfection. In MTT experiments, 1×10^4^ cells of each sample were seeded in each culture well with 150 μL medium containing MTT (0.5 mg/ml).

## Acknowledgments

We appreciate the help equally from the people listed below. They are Professor Yanqiang Liu, Guoqing Liu and Dawei Huang, Associate Professor Bingjun He and Qiang Zhao from College of Life Sciences, Nankai University.

